# Defining the familial fold of the vicilin-buried peptide family

**DOI:** 10.1101/2020.05.26.118075

**Authors:** Colton D. Payne, Grishma Vadlamani, Mark F. Fisher, Jingjing Zhang, Richard J. Clark, Joshua S. Mylne, K. Johan Rosengren

**Affiliations:** The University of Queensland, Faculty of Medicine, School of Biomedical Sciences, St Lucia, Queensland 4069, Australia; The University of Western Australia, School of Molecular Sciences & The ARC Centre of Excellence in Plant Energy Biology, 35 Stirling Highway, Crawley, Perth 6009, Australia; The ARC Centre of Excellence in Plant Energy Biology, 35 Stirling Highway, Crawley, Perth 6009, Australia

## Abstract

Plants and their seeds have been shown to be a rich source of cystine-stabilized peptides. Recently a new family of plant seed peptides whose sequences are buried within precursors for seed storage vicilins was identified. Members of this Vicilin Buried Peptide (VBP) family are found in distantly related plant species including the monocot date palm, as well as dicotyledonous species like pumpkin and sesame. Genetic evidence for their widespread occurrence indicates that they are of ancient origin. Limited structural studies have been conducted on VBP family members, but two members have been shown to adopt a helical hairpin fold. We here present an extensive characterization of VBPs using solution NMR spectroscopy, to better understand their structural features. Four peptides were produced by solid phase peptide synthesis and shown to adopt a helix-loop-helix hairpin fold, as a result of the I-IV/II-III ladder-like connectivity of their disulfide bonds. Inter-helix interactions, including hydrophobic contacts and salt bridges, are critical for the fold stability and control the angle at which the anti-parallel α-helices interface. Activities reported for VBPs include trypsin inhibitory activity and inhibition of ribosomal function, however their diverse structural features despite a common fold suggest additional bioactivities yet to be revealed are likely.

---

Diverse peptide families are found throughout nature and produced by diverse biosynthetic routes.^1–4^ An unusual biosynthetic mechanism, which has been discovered recently, is for a peptide to be proteolytically matured from within a ‘host’ protein that serves an altogether different function.^5^ For these ‘buried’ peptides, a precursor protein is matured into its ancestral protein, as well as an additional peptide that has evolved within a latent region. Two different types of peptides that arise via this biosynthesis in plant seeds have been discovered to date, and both rely on the cysteine-protease, asparaginyl endopeptidase (AEP) for the proteolytic maturation.^6–8^

The first example of this biosynthesis was discovered in sunflowers where the gene *Preproalbumin with SFTI-1* (*PawS1*) encodes a precursor protein for a napin-type seed storage albumin, but also a small cyclic peptide that inhibits trypsin.^5, 9^ A mature seed storage albumin is often a heterodimer consisting of a small and large subunit held together by disulfide bonds. It is during the proteolytic processing of the precursor into the heterodimeric albumin form that the peptide Sunflower Trypsin Inhibitor-1 (SFTI-1) is released.^5^ Albumins are often rich in Cys, Met and Gln residues and are used as a nutrient source during germination by the seedling.^10–11^ Later, other *PawS1* genes were discovered via PCR6 and *de novo* transcriptomics.^12^ This led to the definition of a new family of disulfide containing and usually backbone cyclic peptides called PawS-Derived Peptides and their ancestral relatives, the smaller Cys-less, PawL-Derived Peptides.^6,7^

The existence and extent of a second family of buried plant peptides has now been realized. In 1999, two independent studies reported genes encoding peptides rich in Cys residues buried within precursors of vicilin proteins, another class of seed storage protein. These genes were found in *Cucurbita maxima* (pumpkin)^13^ and *Macadamia integrifolia*.^14^ The encoded preprovicilin protein (PV100) of the pumpkin seed was found to contain the peptide “C2”, buried within the N-terminal region of the precursor sequence.^13^ Interestingly, this peptide is excised from the preprovicilin by AEPs, akin to how SFTI-1 is cleaved from within PawS1. The processing of PV100 results in the mature 50 kDa vicilin, three cytotoxic peptides 4-5 kDa in length and rich in Arg/Glu residues, as well as the 5 kDa peptide C2.^13^ The sequence of the C2 peptide contains four Cys residues, which form a pair of CXXXC motifs connected via disulfide bonds in an I-IV/II-III configuration.^13^ Shortly after C2 was discovered, another group isolated three peptides similar to C2 from macadamia nut kernels.^14^ These peptides, named *Macadamia integrifolia* Antimicrobial Protein 2b-d (MiAMP2b-d), display antimicrobial properties.14 As was the case with C2, the sequences of these three peptides are processed from the N-terminal region of a preprovicilin and contain a pair of CXXXC motifs, however no mechanism for this processing was proposed at the time.^14^ C2 and the three MiAMP2 peptides outlined the potential for a novel buried peptide family, in this case, having evolved from within the precursors of vicilins. Luffin P1, yet another peptide highly similar to C2 and the MiAMPs, was later extracted from the seeds of *Luffa aegyptica*, and shown to be an inhibitor of protein synthesis, but no connection to vicilins was made.^15^

Recent work tied the origin of all these peptides together and showed that they are just the first identified members of a large and ancient family (Figure 1). Zhang *et al.*^8^ established that the precursor of Luffin P1 is indeed a preprovicilin and also showed that precursors with insertions in their N-terminal regions were likely to be the source of many similar hairpin peptides. Luffin P1, C2, MiAMPs, along with many new additions served to establish the Vicilin Buried Peptide (VBP) family.^8^ By confirming that the VBPs are all likely to be AEP-processed, this work also showed that the true sequence of Luffin P1 is longer than originally described.

**Figure 1.**
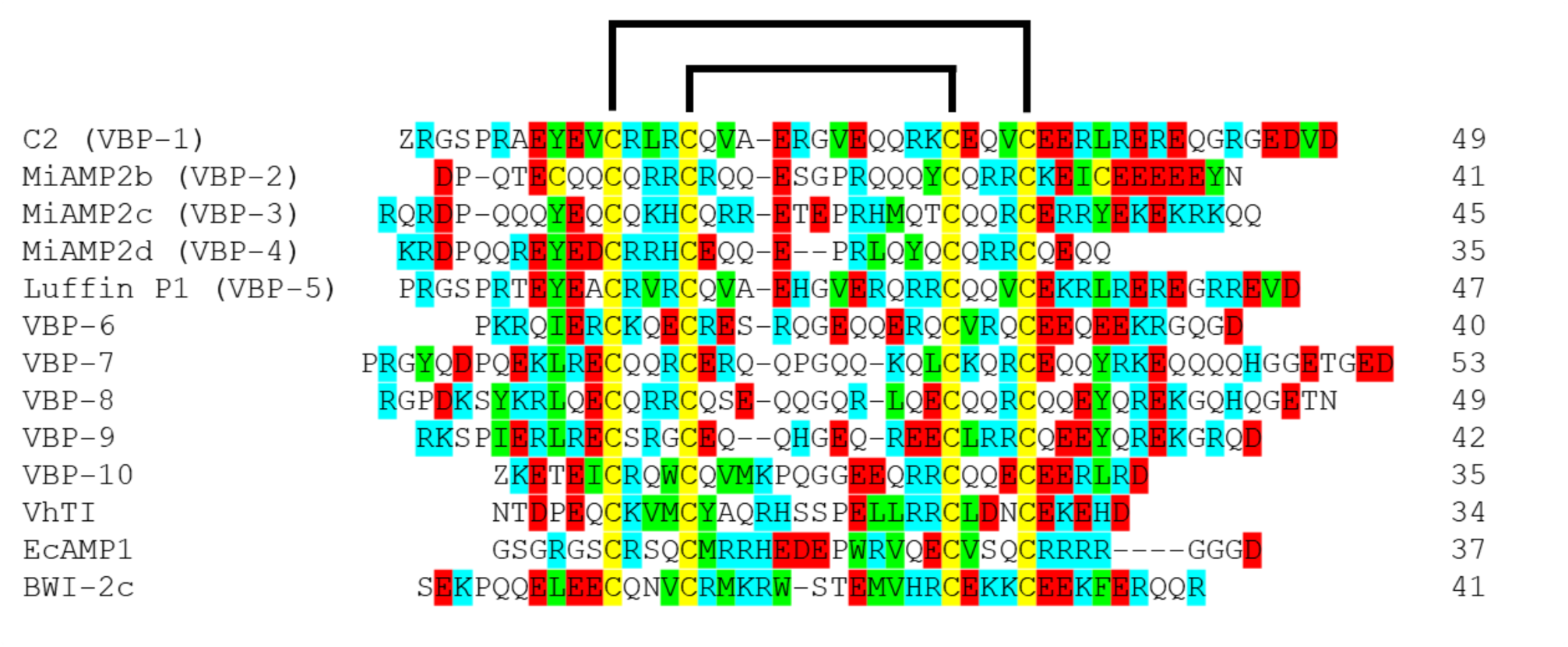
A Sequence alignment of confirmed and putative VBPs, which have been confirmed on a protein level *in planta*. Names, sequences and sequence length. The Cys residues of the CXXXC motifs are highlighted in yellow. Negatively charged residues (Glu and Asp) and positively charged residues (Arg, Lys and His) are highlighted in red and cyan, respectively. Hydrophobic residues are highlighted in green. Sequence alignment produced by Clustal Omega.

Structural studies on this longer Luffin P1 and another VBP from tomato seeds (*Solanum lycopersicum*, VBP-8) showed that they adopt a helix-loop-helix hairpin, which is stapled by two disulfide bonds in a ladder like I-IV/II-III confirmation.^8^ A similar fold has been described for three other bioactive peptides from seeds, *Veronica hederifolia* trypsin inhibitor (VhTI),^16^ *Echinochloa crus-galli* antimicrobial peptide 1 (EcAMP1)^17^ and a buckwheat trypsin inhibitor (BWI-2c).^18^ This suggests that these peptides might also originate from vicilin precursors and that there are potentially other examples of *bona fide* VBPs that have been reported in the literature. In this work we have synthesized and characterized four new peptides to confirm the familial structure and determine the common features essential for folding. We present the full three-dimensional structures of two new VBPs, including the prototypic VBP, C2 from the seeds of pumpkin,13 and detail the structural features of this family.

## RESULTS AND DISCUSSION

### Chemical Synthesis

Recently, a number of sequences have been identified from within a widespread selection of plant seeds and attributed to a new family of peptides known as the vicilin buried peptide family.^8^ To further our understanding of the VBP family, a set of peptides varying in sequence were chosen for structural and functional studies. These included the prototypic C2, VBP-6, and VBP-10. In addition, we chose to make VhTI.^16^ The genetic origin of VhTI has not been confirmed, but its presence in seed, coupled with similarities in size, amino acid composition and disulfide array suggest it is likely to also be produced through processing of a vicilin precursor. The structure of VhTI has been determined by X-ray crystallography in complex with trypsin, but not in free form, so we were interested to see whether this peptide retains its structure in solution. Peptides were assembled in full on resin using Fmoc-based solid phase peptide synthesis (SPPS). After cleavage, the disulfide bonds were randomly or regioselectively formed to match the I-IV/II-III pattern of Luffin P1, VBP-8 and VhTI.^8,16^ Peptides were purified to >95% purity using reverse phase high performance liquid chromatography (RP-HPLC) and their identity confirmed by electrospray ionization mass spectrometry (ESI-MS). MS and HPLC data of all peptides are included as **Figures S1 & S2**.

### NMR Spectroscopy

The only known 3D structures of confirmed VBPs are those of Luffin P1 and VBP-8, which have been determined using solution NMR spectroscopy.^8^ To further expand our knowledge of the structural features of this family and to identify key folding requirements of the family, C2, VBP-6, VBP-10 and VhTI were subjected to homonuclear and heteronuclear NMR spectroscopy. As can be seen in **Figure S3**, the spectral quality of the studied VBPs vary, with differences in levels of signal dispersion, line widths and signal-to-noise ratio being observed. The C2 sample generated the highest clarity spectra with good signal dispersion and sharp, well-resolved peaks for almost all HN signals. This allowed for the full assignment of the HN region of the peptide and almost all other ^1^H signals. Both VhTI and VBP-10 samples generated 1D spectra with good signal dispersion, but poorer signal-to-noise ratio compared to C2. This created some challenges when trying to assign some of the weaker ^1^H signals, including key NOEs in the 2D Nuclear Overhauser Effect SpectroscopY (NOESY) spectra. Despite this, the spectra of C2, VhTI and VBP-10 allowed for the sequential assignment of all backbone signals, as well as the majority of the side chain signals. VBPs are typically rich in Glu, Gln and Arg residues, which results in significant overlap of Hβ and Hγ signals in most spectra. However, distinct signals were observed for these particular residues when located within highly ordered helical regions, allowing for the assignment of these residues with high confidence. A typical sequential walk diagram, that of C2, is given in **Figure S4**. The spectra of VBP-6 differed greatly from the other analyzed VBPs; although maintaining a good signal-to-noise ratio and with sharp signals, VBP-6 lacked significant signal dispersion. This was particularly problematic in the HN region where the overlap of amide signals caused difficulties assigning the backbone chain of the peptide. Ultimately, the combination of overlapping resonances as well as the high density of particular residues such as Glu and Arg, with no unique shifts, prevented assignments of the Hα resonances of residues 1-3, 13, 17 and 21-22. In addition to the homonuclear data, natural abundance ^1^H-^13^C and ^1^H-^15^N HSQC data were analyzed and assigned for C2, however it was only possible to assign the ^1^H-^13^C data for VBP-10 due to the low signal intensity of the ^1^H-^15^N spectra. The spectral assignment of the heteronuclear data was based on the ^1^H chemical shifts derived from homonuclear Total Correlation SpectroscopY (TOCSY) and NOESY data, and allowed for the confirmation of some of the ambiguous ^1^H chemical shifts.

The chemical shifts of ^1^Hα protons are sensitive to backbone torsion angles and analysis of these shifts can help identify secondary structure.^19^ The chemical shifts of all ^1^Hα nuclei were compared to their random coil values and the secondary shifts are presented in Figure 2. Values that deviate susbtantially from random coil are consistent with an ordered structure whereas values closer to zero indicate a disordered random coil-like structure.^20^ From Figure 2 it is clear that C2, VBP-10 and VhTI adopt highly ordered structures, but VBP-6 appears to have limited secondary structure and mainly adopts random coil. The stretches of negative secondary Hα shifts seen in C2, VBP-10 and VhTI are consistent with two helical regions separated by a turn. Given the presence of the two disulfide bonds between the helical regions it could be predicted that the ordered VBPs are adopting a helix-loop-helix fold, like Luffin P1, VBP-8 and VhTI, the last when in complex with trypsin.

**Figure 2.**
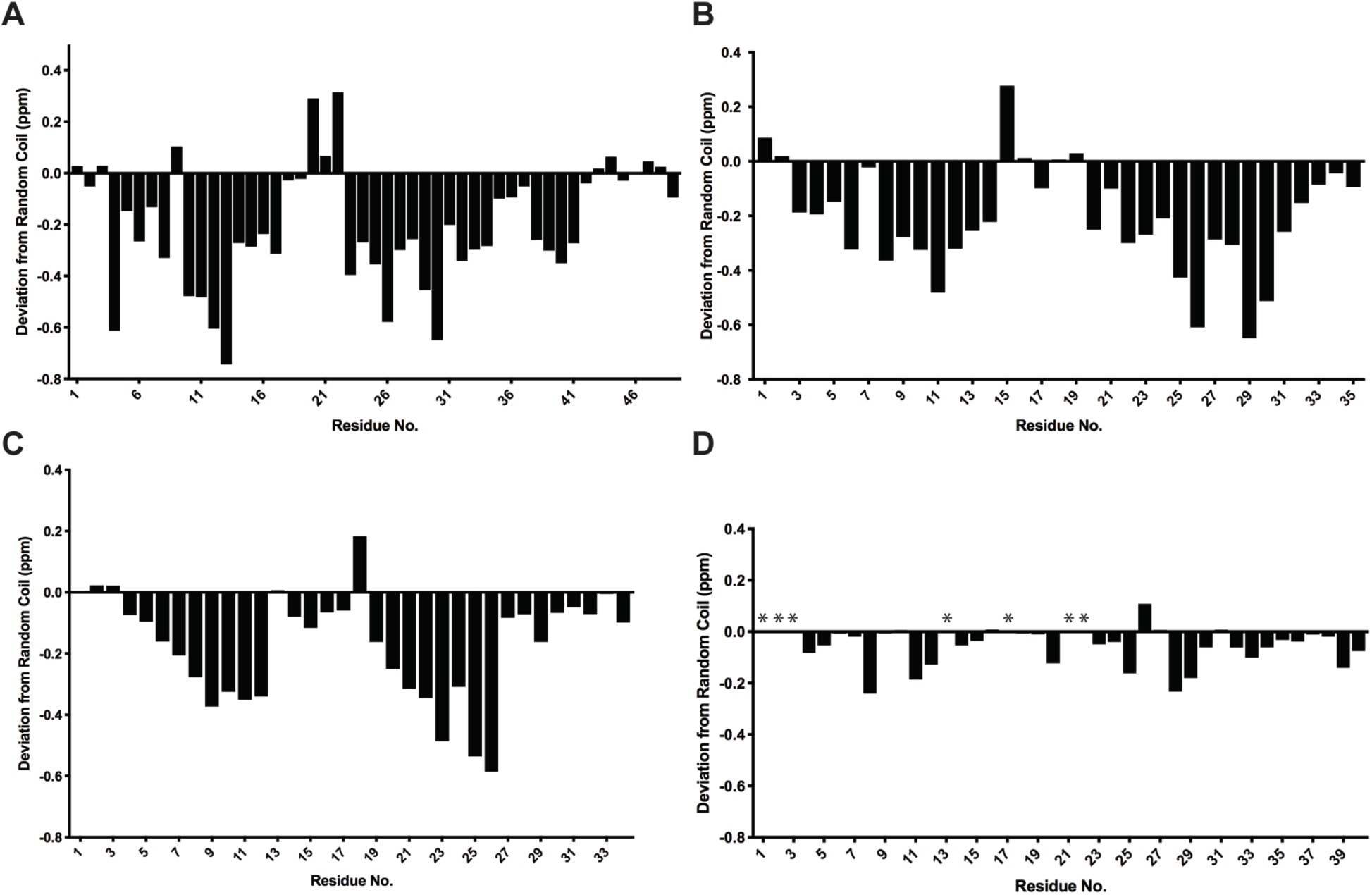
Secondary Hα shifts of synthetic VBPs. (A) C2; (B) VBP-10; (C) VhTI; and (D) VBP-6. Observed significant negative stretches are indicative of helical regions, with the positive or close to zero values in the center of these negative stretches being consistent with the loop region of the hairpin.

### Structure Determination of C2 and VBP-10

To further evaluate the structural features of VBPs the full 3D structures were determined for C2 and VBP-10 following established protocols.21 For structure calculations, a combination of inter-proton distance restraints derived from NOESY cross peak volumes, hydrogen bond restraints based on temperature coefficients and analysis of preliminary structures, as well as Torsion Angle Likelihood Obtained from Shift and sequence similarity (TALOS-N) defined backbone dihedral angles were used. Preliminary structure calculations were performed using automated NOE assignment within CYANA, with the final structure calculation and water minimization performed in CNS. The structures were analyzed using MolProbity to determine the quality of the structural geometry and atom packing. The 20 best structures from the final 50, based on MolProbity scores, low energy, and no significant experimental violations were chosen to represent the solution structures of the each peptide. These structures are shown as both superposed backbone ensembles and in ribbon representation in Figure 3. The statistics relating to these structures are listed in Table 1. There are a notably reduced number of inter-proton distance restraints present in VBP-10 compared to C2, in particular no NOEs could be unambiguously assigned as long range across the helices. Resonances in VBP-10 were generally broader and weaker, which may suggest that a more dynamic helical interface is the reason for the lack of such NOEs, but it could also be the result of resonance overlap of key side-chain to side-chain NOEs. Both C2 and VBP-10 have good stereochemical quality, with minimal Ramachandran outliers. These data highlight that both peptides have well-defined helical regions with backbone RMSD values below 0.65 Å. The structures generated for both C2 and VBP-10 were above the 75th percentile of all structures according to MolProbity.

**Table 1.** NMR Structural Statistics of C2 and VBP-10.

| Energies (kcal/mol) | C2 | VBP-10 |
| --- | --- | --- |
| Overall | -1831.4 ± 92.2 | -1349.7 ± 75.2 |
| Bonds | 14.5 ± 0.95 | 9.7 ± 1.0 |
| Angles | 57.8 ± 5.0 | 40.2 ± 4.0 |
| Improper | 22.0 ± 2.7 | 13.1 ± 2.5 |
| Dihedral | 221.0 ± 1.7 | 162.3 ± 2.4 |
| Van der Waals | -190.3 ± 9.9 | -122.7 ± 5.9 |
| Electrostatic | -1956.9 ± 95.5 | -1452.8 ± 78.7 |
| NOE | 0.043 ± 0.025 | 0.041 ± 0.011 |
| cDih | 0.31 ± 0.19 | 0.46 ± 0.26 |
| MolProbity Statistics |  |  |
| Clashes (>0.4 Å / 1000 atoms) | 13.9 ± 2.9 | 7.1 ± 3.0 |
| Poor rotamers | 0.05 ± 0.22 | 0.15 ± 0.37 |
| Ramachandran Outliers (%) | 0.54 ± 0.96 | 1.56 ± 2.38 |
| Ramachandran Favored (%) | 95.4 ± 2.3 | 93.0 ± 3.6 |
| MolProbity score | 1.95 ± 0.18 | 1.82 ± 0.21 |
| MolProbity score percentile | 77.7 ± 8.4 | 83.5 ± 8.7 |
| Atomic RMSD (Å) |  |  |
| Mean global backbone (of helical regions) | 0.63 ± 0.18 | 0.59 ± 0.17 |
| Mean global heavy (of helical regions) | 1.65 ± 0.21 | 1.96 ± 0.22 |
| Experimental Restraints |  |  |
| Distance restraints |  |  |
| Short range (i-j < 2) | 348 | 207 |
| Medium range (/i-j/ < 5) | 30 | 20 |
| Long range (/i-j/ > 5) | 12 | 0 |
| Hydrogen bond restraints | 36 (18 H-bonds) | 40 (20 H-bonds) |
| Total | 402 | 267 |
| Dihedral angle restraints |  |  |
| φ | 35 | 25 |
| ψ | 37 | 26 |
| χ<sup>1</sup> | 4 | 4 |
| χ<sup>2</sup> | 0 | 3 |
| Total | 76 | 58 |
| Violations from experimental restraints |  |  |
| Total NOE violations exceeding 0.2 Å | 0 | 0 |
| Total dihedral violations exceeding 2.0° | 0 | 0 |

**Figure 3.**
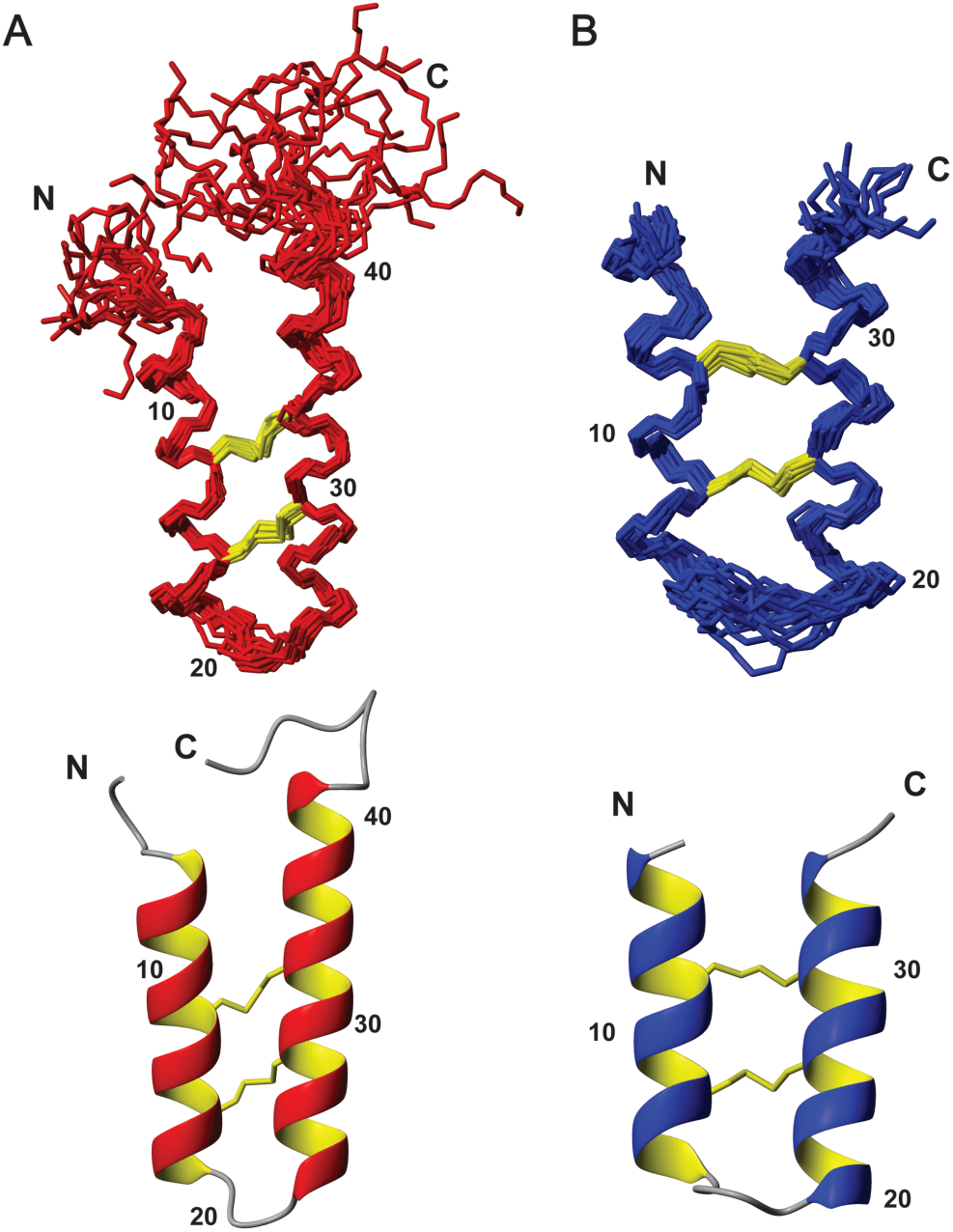
Solution NMR structures of C2 (A) and VBP-10 (B). Top images show the structural ensembles in stick format, whereas the lower panels show the lowest energy structures in ribbon format. Disulfide bonds that crosslink the α-helices and form the hairpin structure are shown in yellow stick format, with selected residue number labels supplied for orientation.

### Structural Features of C2 and VBP-10

Both C2 and VBP-10 adopt helix-loop-helix structures that are similar to the previously reported VBPs.^8^ The N-terminal helices are shorter than the C-terminal, 15 vs. 20 and 11 vs. 14 residues in C2 and VBP-10, respectively. The fold is primarily stabilized by the ladder like I-IV/II-III disulfide bond connectivity and a large number of backbone hydrogen bonds in the i-i_+4_ pattern characteristic of α-helices. The conformation and positioning of the disulfide bonds differ, with VBP-10 having a longer central loop, while C2 has significant sequence extensions both at the N-terminus (5 residues) and the C-terminus (10 residues). Notably, the spatial arrangement differs with C2 showing a close to planar alignment of its anti-parallel helices, while in VBP-10 the anti-parallel helices vary greatly in alignment demonstrating angles of 7° to 34° between the C-terminal helix and the N-terminal helix. This is related to sequence differences generating different packing of side chains at the inter-helical interface. Clearly defining the precise tilt angle, if a preferred value exists, in VBP-10 is impossible without inter-helical NOEs. In C2, a notable hydrophobic interaction, which is supported by a number of NOE contacts, results from the packing of Tyr9 and Leu37. The positioning of oppositely charged residues also results in several potential surface exposed salt-bridge interactions stabilizing the fold, including between Glu8 and Arg36, as well as Arg13 and Glu30. VBP-10 contains a single aromatic residue, Trp10, but this is in contrast to Tyr9 in C2 positioned on the outside of the helical face and its side-chain resonances do not show any intra- or inter-helical NOEs. Like C2, VBP-10 does contain several potential salt-bridge interactions, including between Glu3 and Arg32 as well as Lys15 and Glu21.

### Comparison of C2 and VBP-10 to other VBPs

The combination of sequence and structural data now allows insight into the conserved structural features of the VBP family. The sequence alignment in Figure 1 highlights the positions of charged and hydrophobic residues. The CXXXC motifs highlighted in yellow are the only absolutely conserved feature of all members of the VBP family, allowing for the identification of VBPs present within the N-terminal region of vicilin precursors. VBPs as a whole are primarily comprised of Glu, Arg and Gln residues, with few aromatic and hydrophobic residues present. Despite the limited complexity in the amino acid composition of the VBP sequences, there is low sequence similarity between individual members of the family. Typically between any given VBP there is less than 40% similarity, with the exceptions being C2 and Luffin P1, which are both from cucurbit species and share 72% similarity. Additionally, VBP-10 shows 56% and 59% similarity to C2 and Luffin P1, respectively.

Apart from the Cys residues, a pair of hydrophobic residues at the positions three residues before the first Cys and four residues after the last Cys are a conserved feature. These residues are commonly Tyr and Leu (Figure 1). Intriguingly, which of the two positions contains a Leu and which a Tyr residue is interchangeable. When present, these Tyr and Leu residues always end up in close proximity and interacting because of the helical hairpin, indicating the importance of these residues for the structural stabilization of the fold (Figure 4). The interaction is easily confirmed by the presence of multiple strong NOEs between the Tyr aromatic protons and the Leu methyl protons, as noted above for C2. In another putative member of the family, BWI-2c, the hydrophobic pair is Leu-Phe rather than Leu-Tyr, and NOEs are observed also between these residues.^18^ This type of interaction, although clearly favored, is however not essential. In MiAMP2b these two positions are instead occupied by additional cysteine residues, which presumably form a third disulfide bond stabilizing the hairpin via a third covalent crosslink rather than a hydrophobic interaction. We show here that VhTI and VBP-10 are still able to adopt the same overall fold without notable interactions between these positions, and the other putative VBP, EcAMP, has previously been shown to adopt the helix-loop-helix fold despite containing Arg at both positions. VhTI still contains multiple Leu residues and a singular Tyr residue within its sequence. Despite these residues not being in the same location as other VBPs, the NOESY data suggest that two of the Leu residues and the Tyr residue are in close proximity and also participate in inter-helix hydrophobic interactions. This is consistent with what is seen in the crystal structure of VhTI bound to trypsin.^16^ Similarly, in EcAMP1 Met12, Pro19, Val22 and Val26 form a hydrophobic cluster that packs up against the disulfide core.^17^ VBP-10 by contrast does not contain any significant hydrophobic interactions across its helices, which might explain the lack of a defined arrangement and NOEs.

**Figure 4.**
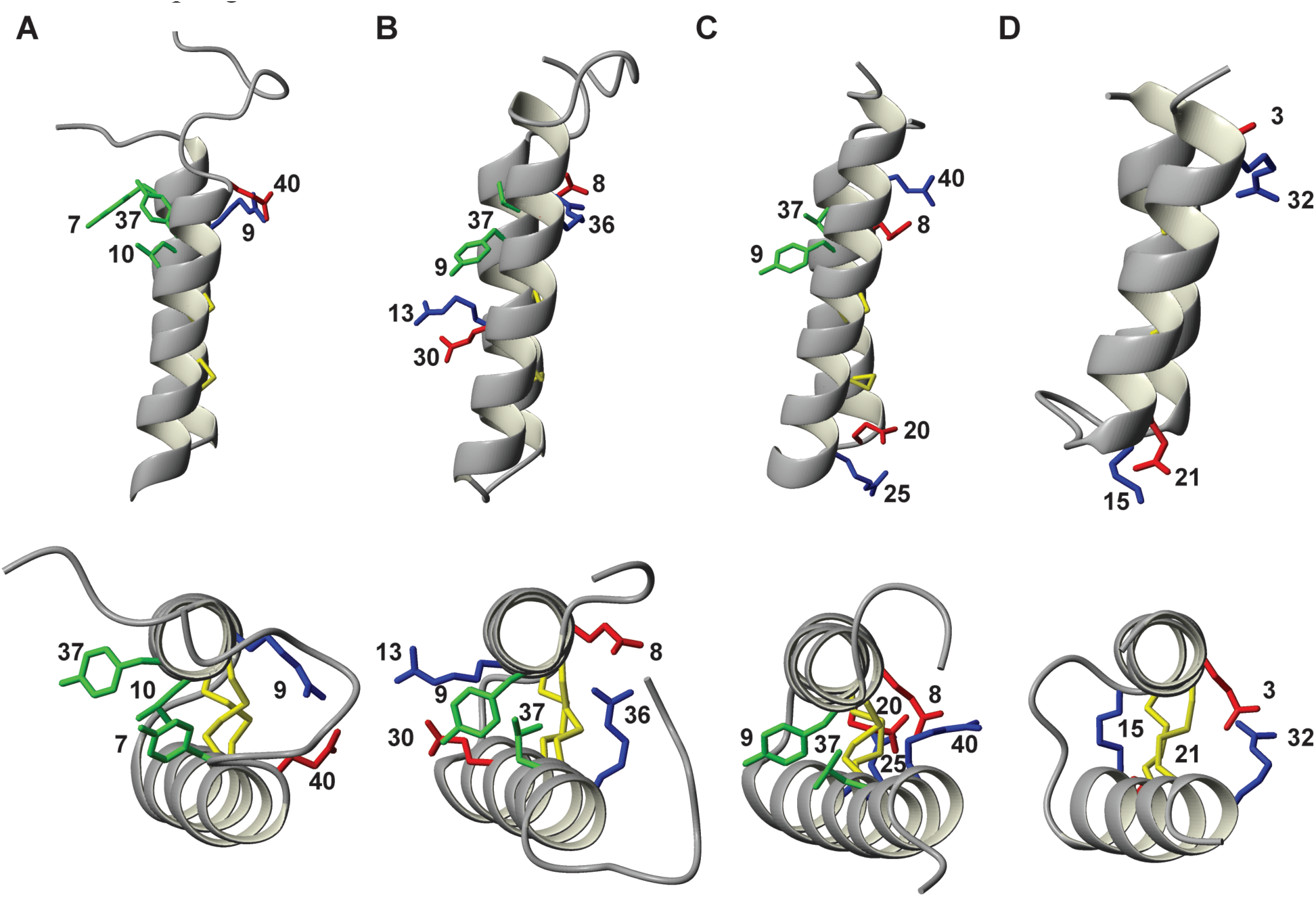
Structural comparison presenting side-on (upper panels) and top-down (lower panels) views of confirmed VBPs in ribbon format. (A) VBP-8;^8^ (B) C2; (C) Luffin P1^8^ and (D) VBP-10. The N-terminal helix is at the back of the image in the side-on model and on top in the top-down model and inter-helical interactions are highlighted. Positively charged Arg and Lys residues are colored blue, negatively charged residues Glu and Asp are colored red, hydrophobic Leu and Tyr residues participating in hydrophobic interactions are colored green and disulfide bonds colored yellow. Side chains are labeled with residue numbers.

In addition to local hydrophobic interactions, ionic interactions are also commonly observed between positively and negatively charged residues within VBP structures. Commonly, these salt bridges are intra-helical, stabilizing the secondary structure, however inter-helical ionic interactions also stabilize the tertiary interface of the α-helices. All VBPs that have been structurally studied contain multiple intra-helical salt bridges on the external faces of the α-helices and additionally contain at least one inter-helical salt bridge (Figure 4). VBP-10 contains two, one at either end of the helical segments, and these interactions appear sufficient for the overall fold in the absence of hydrophobic interaction. The location of the salt bridges in VBPs are non-uniform, being observed between residues close to the termini of the VBP, within the two CXXXC motifs and in the loop region.

From both the secondary chemical shifts and the calculated structures it is clear that the length of the loop separating the helices is of variable length, ranging between two and five residues. VBP-8 has previously been reported to have a short two-residue loop,8 with VBP-10 from this work having a longer five-residue loop. This difference in loop length allows for increased flexibility in the region and in particular allows different interfacing angles of the α-helices. VBP-8 contains the shortest loop of all VBPs and has the smallest interface angle of the VBPs (Figure 4). In contrast, VBPs such as Luffin P1 and VBP-10 have larger interface angles and as a result of the helical termini ending up further apart, require larger loop regions. These VBPs also appear to have more extensive inter-helical interactions occurring at both the terminal and loop ends of the α-helices.

### Why does VBP-6 not adopt a helix-turn-helix fold?

VBP-6, unlike all other studied VBPs, does not adopt an ordered structure in solution, evident from largely random coil chemical shifts. VBP-6 lacks hydrophobic residues in positions where they can interact with each other in a folded state. Like all VBPs, VBP-6 does contain a multitude of charged residues, primarily Arg, Lys and Glu. However, a closer analysis of their position suggests that the spacing of these residues is unfavorable for a folded state, in contrast to VBP-10, which is able to retain a helix-turn-helix fold despite a lack of hydrophobic packing. Figure 5 shows a helical wheel diagram for VBP-10 and what the equivalent schematic view for VBP-6 would have looked like if it had adopted a helix-loop-helix fold. From this diagram it can be seen that the only possible inter-helical salt bridge that could be formed in VBP-6 would be between Lys9 or Arg16 and Glu22, as indicated by the green dashed lines. These residues are on the loop side of the disulfide array rather than the terminal part of the fold, and may not offer sufficient stabilization. More importantly, the position of charged residues within the putative helical segments are not ideal in VBP-6. In particular the C-terminal helix has a high density of negative charges, including Glu30, Glu31, Glu33 and Glu44 that would be expected to clash in a helical conformation. Potential favorable and unfavourable ionic interactions between residues which are three or four residues apart within the helices are highlighted in Figure 5. In VBP-10 there are seven potentially favorable interactions and only one that is potentially unfavorable. In contrast in VBP-6 eight favorable interactions are cancelled out by eight potentially unfavorable ones. This probably prevents this peptide from adopting the familial VBP fold.

**Figure 5.**
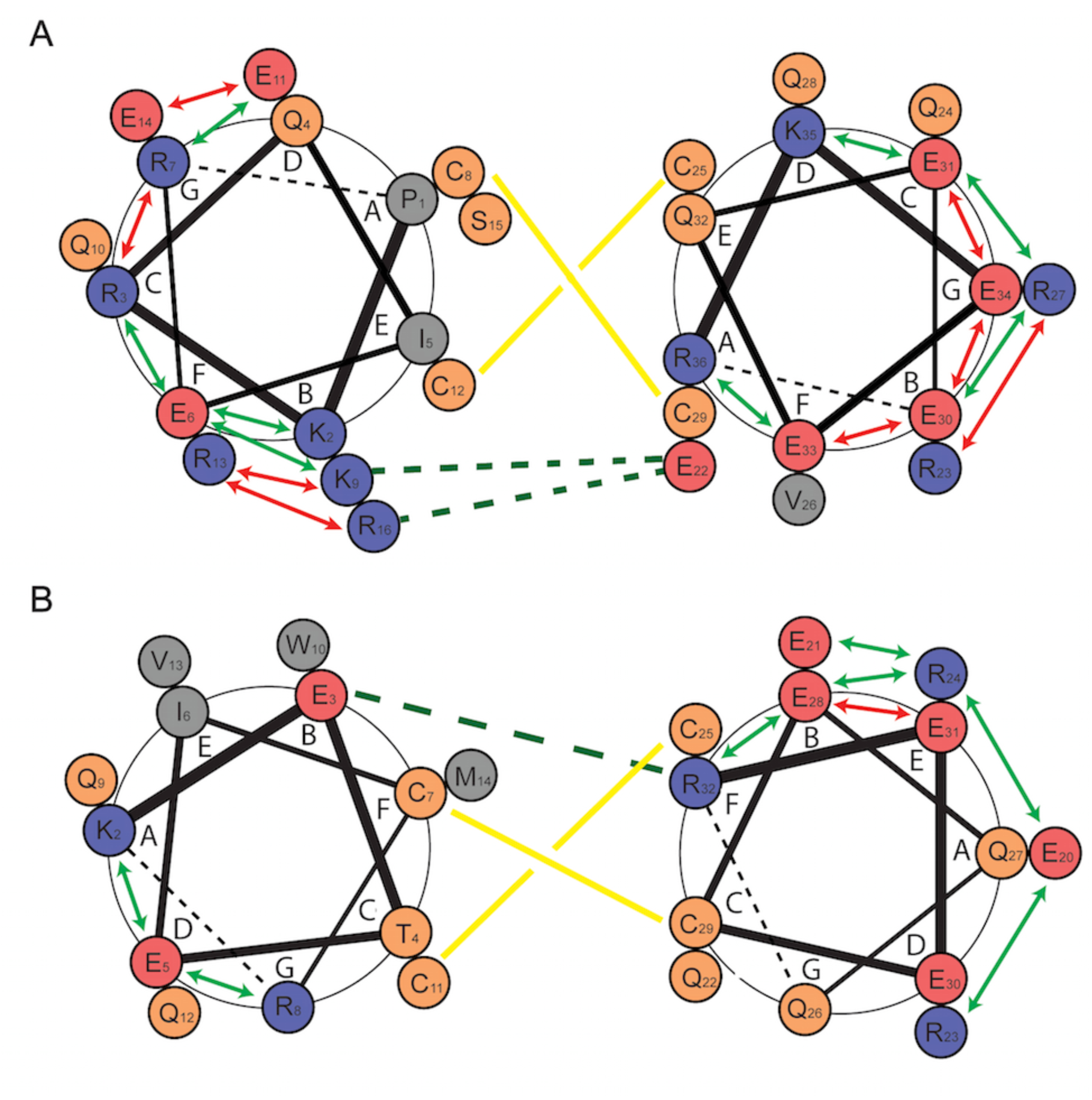
Schematic comparison of ionic interactions within the helices of VBP-6 and VBP-10. (A) Putative helical wheel diagram for VBP-6. The left N-terminal helix comprises residues Pro1 to Gln17 and the right C-terminal helix comprises residues Glu20 to Arg36, illustrating a minimal two-residue loop gap. (B) Helical wheel based on the structure of VBP-10. The N-terminal residues of the helices are located at position A of the wheel, with subsequent residues in sequence at positions B, C, D, E, F and G. In the N-terminal helix lines go from thick to thin as the helix progress away from the viewer, whereas in the C-terminal helix the connecting lines go from thin to thick as the helix progress towards the viewer. Residues are labeled with residue numbers. Positively charged residues are indicated with blue circles, negative with red circles, polar residues with orange circles and hydrophobic residues with gray circles. Yellow lines represent the disulfide bonds, with the solid yellow line being the I-IV disulfide bond and the broken yellow line being the II-III disulfide bond. The green dashed lines represent potential inter-helical salt bridges. Red and green arrows highlight potentially favorable and unfavorable interactions between residues three or four residues apart within the helical segments, respectively.

### Comparison to other helix-loop-helix peptides

VBPs are far from the only family of peptides that adopt a helix-loop-helix hairpin fold. This fold is adopted by a wide range of peptide sequences from different sources including numerous sea creatures^22–24^ and scorpions,^25–26^ and also by plant peptides that are not members of the VBP family.^17–18^ Peptides adopting this fold can greatly vary in size, from as short as 23 residues in the case of the Om-toxins of the *Opsithacanthus madagascariensis* scorpions25 to as large as 55 residues in the case of neurotoxin B-IV from *Cerebratulus lacteus* marine worm.^22^ The disulfide bond configuration of these peptides is maintained in all observed cases, that is, a ladder-like configuration. However, the number of disulfide bonds present is 0, 2 or 4,^18, 22–23^ with no observed helix-loop-helix fold structurally described to date containing 1 or 3 disulfide bonds. These peptides have a wide variety of functions both defensive, as in the case of antimicrobial plant peptides^17^ and neurotoxic excretions of the Red Sea Moses sole flatfish,^27^ and offensive as in the case of neurotoxic venom of cone snails^24^ and potassium channel blockers of scorpions.^25^ Despite variations in function, origin and disulfide bond number, the overall structures of these peptides are highly similar, as can be seen in Figure 6. The similarities extend past the basic fold, with features such as inter-helical hydrophobic interactions being analogous to the VBPs. Tyr and Leu residues similar to those of the VBP family are seen also in these peptides, however other combinations of hydrophobic residues are also found in similar interactions. Additionally, ionic interactions between charged residues are also commonly observed as inter-helical interactions, again similar to those observed in the VBP structures. Thus the VBPs largely conform to rules that are not specific to this family but common to the general helix-loop-helix fold.

**Figure 6.**
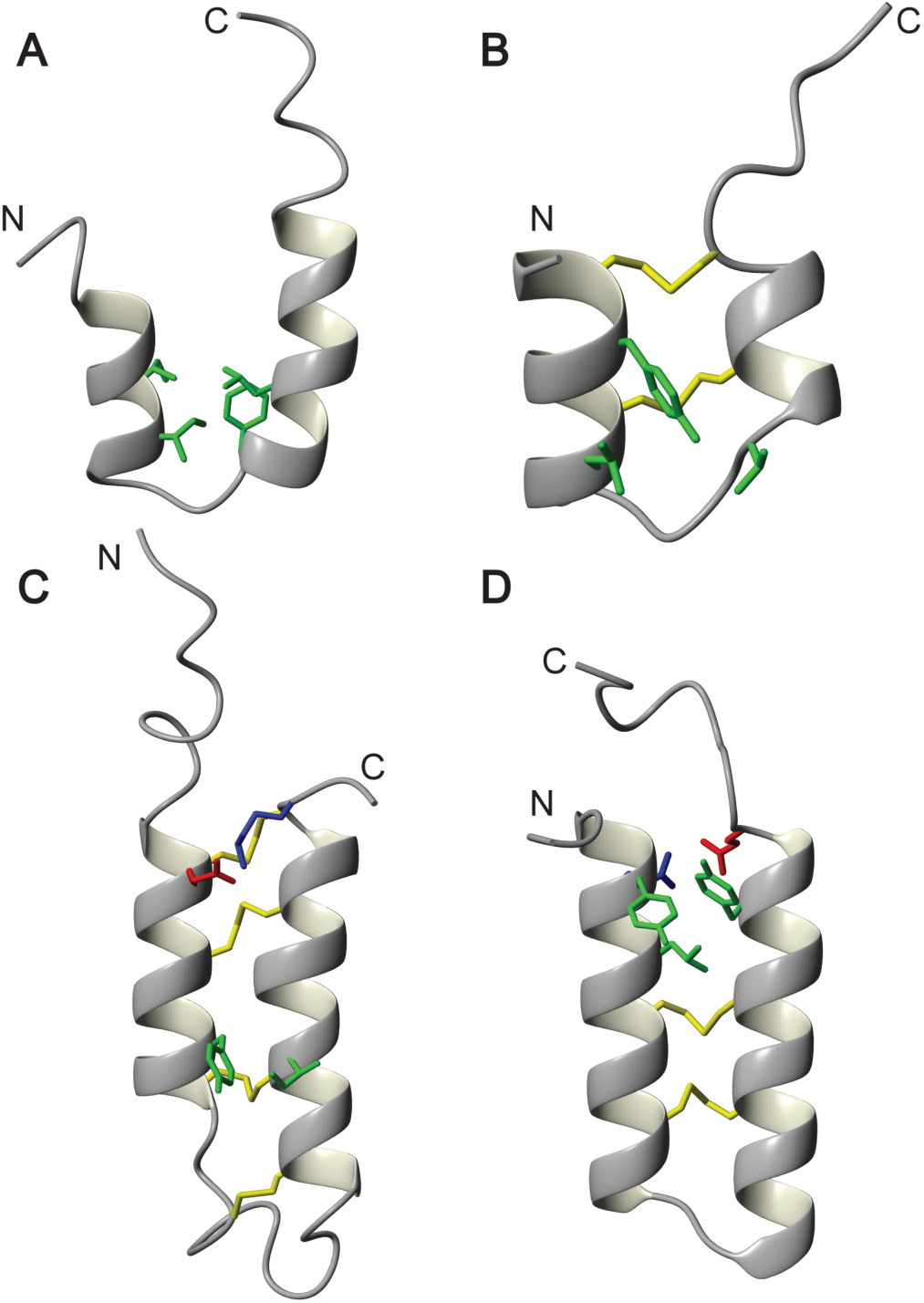
Structural comparison of helix-loop-helix peptide from other sources. (A) Antimicrobial peptide Pardaxin Pa4 from the Red Sea Moses sole fish *Pardachirus marmoratus*^23^; (B) Potassium channel blocker OmTx1 from the scorpion *Opisthacanthus madagascariensis*^25^; (C) Neurotoxic peptide B-IV from the marine worm *Cerebratulus lacteus*^22^ and (D) VBP-8^8^. Ribbon representation of the helix-loop-helix fold peptides with the N-terminal helix on the left and the C-terminal helix on the right. Residue side-chains that are participating in notable inter-helical interactions are shown. Positively charged Arg and Lys residues are colored blue, negatively charged residues Glu and Asp are colored red, hydrophobic Leu and Tyr residues are colored green and disulfide bonds colored yellow.

### Trypsin Inhibition by VBPs

Given that C2 as well as the putative VBPs VhTI and BWI-2c have been reported to inhibit trypsin, and indeed the crystal structure of VhTI in complex with trypsin has been solved,^16^ we wanted to investigate whether this is a common bioactivity among these peptides. C2, VhTI, VBP-10, and VBP-6 were all subjected to a trypsin inhibition assay, wherein all peptides as well as a positive control, a Bowman-Birk inhibitor, were incubated at a range of concentrations from 0.01 µM to 8 µM (Figure 7). As expected, VhTI potently inhibits trypsin with an IC50 of ~0.4 µM,^16^ but no intriguinginly inhibition was observed for C2, VBP-10 or VBP-6.

**Figure 7.**
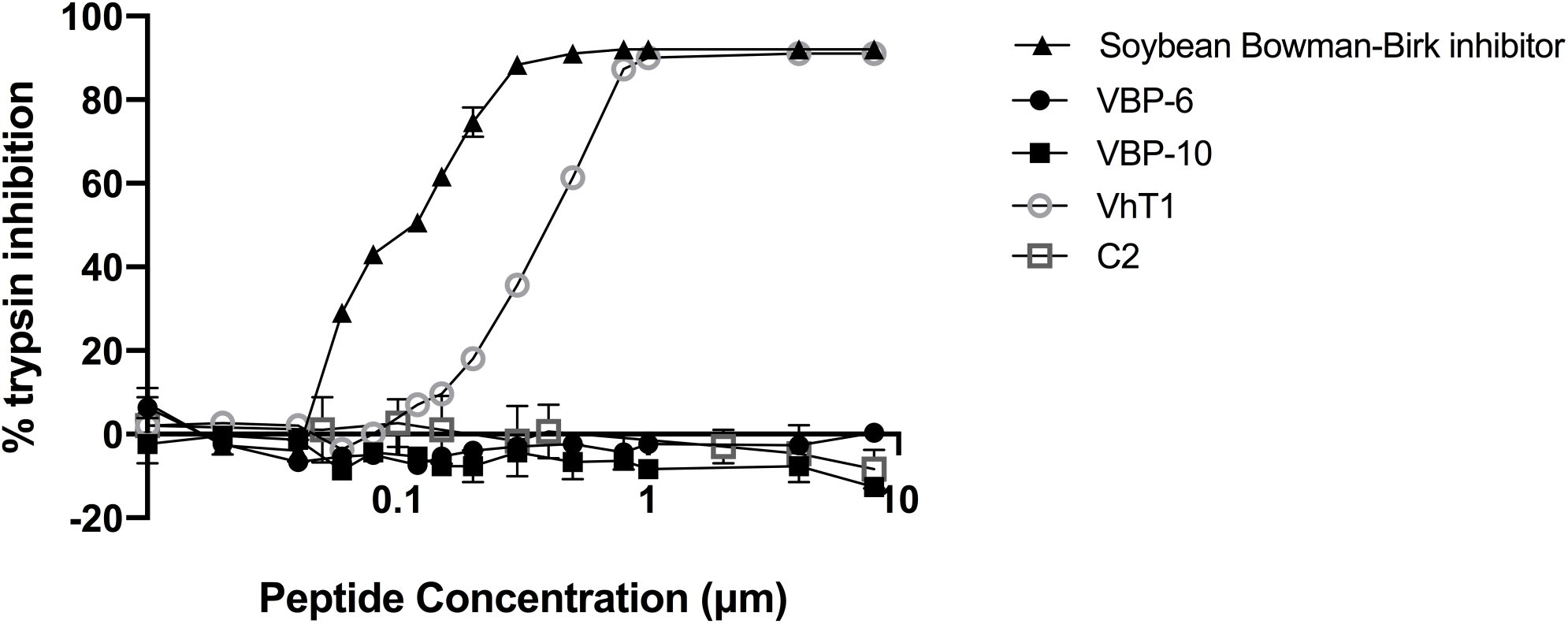
Inhibition of bovine trypsin by VBPs. Trypsin activity was determined in the presence of C2, VhTI, VBP-6 and VBP-10 at a range of concentrations from 0.01 to 8 µM. Soybean Bowman-Birk inhibitor was used as a positive control. The standard error of the mean is shown for each triplicate data point.

A helix-loop-helix fold is an unusual structure for engaging proteases, with most protease inhibitors adopting a beta-strand structure within the protease active site.^28^ VhTI contains an Arg at position 15, four residues after the second cysteine residue. This Arg is capable of inserting itself into the binding pocket of trypsin despite residing within a helical motif.^16^ In addition, Met10, Ala13 and Gln14 form contacts with trypsin. The position of these residues in the bound form of VhTI is compared to the positions of equivalent residues in C2 and BWI-2c in Figure 8. BWI-2c adopts a near identical conformation in solution, projecting the side chains of Val14, Met17, Lys18 and Arg19 in a fashion that would be expected to be able to engage trypsin, and this is consistent with the inhibition assays by Conners *et al.*^16^ In contrast C2, which has also been reported as a trypsin inhibitor, has an Arg five residues after the second cysteine, at position 21, and this residue, based on the structure determined here, would seem unlikely to bind to the P1 pocket of trypsin, consistent with the lack of activity we observe in our work (Figure 7). We have previously reported for the PDP family that even small changes of conformation of side chains can prevent trypsin inhibition in peptides that are highly homologous to potent inhbitors.^29^ It is unclear why activity was observed previously,^13^ but given it was plant-extracted material that was tested (versus our synthetic C2) a bioactive contaminant of plant origin cannot be ruled out. We recently showed that neither Luffin P1 nor VBP-8 is active against trypsin,^8^ and looking at their sequences this was not surprising given that both lack a Lys or Arg at an equivalent position to VhTI in this loop. VBP-10 tested here, on the other hand, does contain Lys15 at a suitable position four residues after the second cystine. The fact that it nonetheless cannot inhibit trypsin highlights the requirement for additional distinct structural features, and that protease inhibition is unlikely to be a general mechanism of action for VBPs. VBP-10 contains a Pro immediately after Lys15 which may prevent the interaction as trypsin cleavage before Pro is not favored. Given the lack of conserved sequences, although trypsin inhibition is possible, it would seem highly unlikely that most VBPs would share any common bioactivity, but rather might have evolved distinct functions.

**Figure 8.**
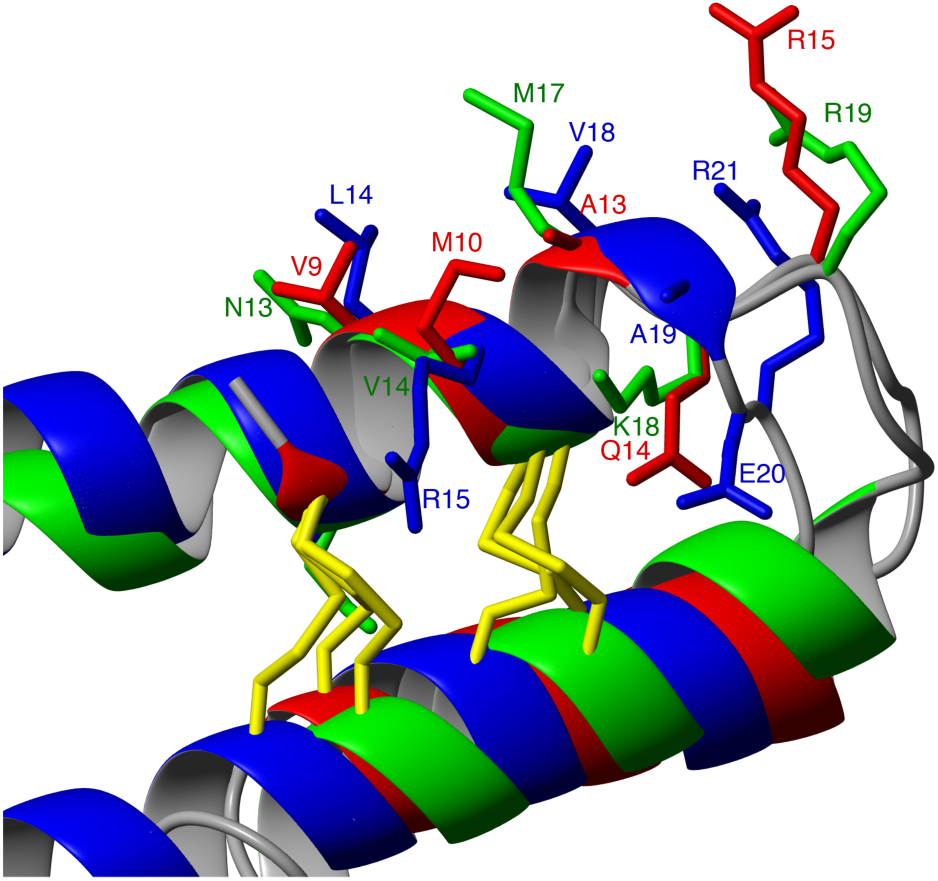
Comparison of structures of VhTI (red), BWI-2c (green) and C2 (blue). Disulfide bonds are shown in yellow. Side chains important for trypsin inhibition in VhTI and the equivalent residues in BWI-2c and C2 are shown in stick format and labelled with residue and number.

### Are VhTI, EcAMP1 and BWI-2c bona fide VBPs?

The realization that other seed peptides, whose genetic origin has not been confirmed, adopt a similar fold to the VBPs suggest that there might be more members of the family already described in the literature. We looked for available transcriptomes from *Veronica hederifolia*, *Echinochloa crus-galli* and *Fagopyrum esculentum* seeds and found that although none is available for the former two, transcriptomic data is available for buckwheat seeds (*Fagopyrum esculentum*), that contain BWI-2c. Using the publicly available data, assembling de novo transcriptomes for *F. esculentum* could only provide a partial precursor sequence for BWI-2c (**Figure S5**). The sequence surrounding the BWI-2c sequence is repetitive and similar to the repeating regions encoding multiple VBPs within Class IV vicilin precursors.^30^ Also present flanking BWI-2c are candidate AEP target sites. As was found for Luffin P1, were BWI-2c to be AEP processed, its sequence might actually be 51-residues starting with Pro and ending with Asp, instead of 41-residues beginning with Ser and ending with Arg. The partial sequence we found also has the potential to encode other VBPs not yet reported in the literature. More work is required to obtain the full precursor sequence for BWI-2c, but in addition to a similar 3D structure, this partial precursor also supports a biosynthetic origin for BWI-2c like other VBPs.

This study sought to investigate the structural features of the VBP family and define its familial fold. Three confirmed and a putative fourth VBP were synthesized using SPPS and studied by solution NMR spectroscopy. VBP-10 and C2 adopted helix-loop-helix folds similar to the one previously reported for VBPs, but a lack of hydrophobic interactions at the helical interface of VBP-10 and broader signals suggest the helices may not be locked into a specific position relative to each other. The trypsin inhibitor peptide VhTI was confirmed to adopt a helix-loop-helix fold in solution based on secondary chemical shifts, which was consistent with its structure bound to trypsin. By contrast, VBP-6 did not adopt an ordered fold, and consequently is the first member of the VBP family to be unstructured. Inter-helical interactions that support the conserved disulfide bonds appear to be essential for the fold. In addition to hydrophobic interactions, salt bridges are common between charged residues, and the type and position of these interactions guide the inter-helical angle. We confirmed the trypsin inhibitory activity of VhTI, and showed that this feature was not conserved in VBP-6 or VBP-10. Surprisingly we were not able to reproduce the reported activity for C2, but its structure does not seem to be congruent with trypsin inhibition. The physiological significance of VBPs remains an enigma, but those with trypsin inhibition probably evolved as a defense against gramnivores. Given the wide spread of bioactivities reported for other helix-loop-helix peptides, different VBPs might contribute to defense through mechanistically unrelated and yet to be discovered activities.

## EXPERIMENTAL SECTION

### Solid Phase Peptide Synthesis

All peptides were synthesized on a 0.125 mmol scale using fluorenylmethyloxycarbonyl (Fmoc)-based solid phase peptide synthesis. The resin used as an anchor for peptide assembly was Tentagel XV 4-hydroxymethyl phenoxyacetic acid (Rapp Polymere, GmbH). Prior to the loading of the C-terminal residue, the resin was swollen in dimethylformamide (DMF) for 24 h. A loading mixture containing the C-terminal residue was prepared by dissolving ten equivalents of amino acid in dichloromethane (DCM) and five equivalents of N,N’-diisopropylcarbodiimide. This solution was stirred for 30 min at 4°C, before removal of the DCM via rotary-evaporation. The then dried amino acid was re-dissolved in the minimum DMF required for complete solvation. This solution was added to the swollen resin with equivalents of 4-dimethylaminopyridine and shaken for 12 h. The entire loading process was repeated to achieve adequate loading. A CS336X automatic peptide synthesizer (CSBio) was used to couple the remaining residues in the peptide chain, with the coupling cocktail being applied to the resin twice for all amino acids containing branched β-carbons. Prior to the coupling of each residue, 20% (v/v) piperidine in DMF was used for the deprotection of the N-terminal Fmoc-group. After deprotection and washing with DMF, each residue was coupled using eight equivalents of amino acid, four equivalents of N,N,Nʹ,Nʹ-tetramethyl-O-(1H-benzotriazol-1-yl)uranium hexafluorophosphate and eight equivalents of N,Nʹ-diisopropylethylamine. Once the peptide chain was completed a final deprotection was conducted before washing with DMF, followed by DCM, and then the resin was dried under nitrogen. Peptide was cleaved from the resin by addition of 50 mL of a cocktail of trifluoroacetic acid, triisopropylsilane, 3,6-dioxa-1,8-octanedithiol and water (95:2:1.5:1.5). Following cleavage the trifluoroacetic acid was reduced via rotary-evaporation and the peptide precipitated from solution using cold diethyl ether. The precipitated peptide was filtered and re-dissolved in a solution of MeCN/water (50:50) before lyophilization.

### Peptide Purification and Folding

Crude peptide was purified via RP-HPLC using a solution of 90% acetonitrile and 0.05% trifluoroacetic acid at a gradient of 1%/min on a preparative C18 column (300 Å, 10 µm, 21.20 mm i.d x 250 mm, Phenomenex). ESI-MS was used to confirm the mass of the peptide. Regio-selective disulfide bond formation using acetamidomethyl and trityl-protected Cys residues was employed to generate the desired connectivity of I-IV/II-III. The formation of the first disulfide bond, between Cys II-III, was conducted using a solution of 0.1 M ammonium carbonate, pH 8.3, at a peptide concentration of 0.25 mg/mL for 24 h. After this further purification was conducted using a semi-preparative C18 column (300 Å, 5 µm, 10 mm i.d. x 250 mm, Vydac). The removal of the acetamidomethyl protecting groups and formation of the second disulfide bond was conducted via iodolysis in acetic acid/water (50:50) at a concentration of 0.25 mg/mL. A solution of 0.1 M iodine was added to the peptide solution until a noticeable color change from clear to orange was achieved. This solution was then stirred under nitrogen in a dark environment for 4 h, prior to quenching with ascorbic acid. Final purification was conducted using the previously mentioned method and purity was determined using a C18 analytical column (300 Å, 5 µm, 2.1 mm i.d. x 150 mm, Vydac).

### NMR Spectroscopy

Samples were prepared by dissolving 0.8-2 mg of peptide in 500 µL of H_2_O/D_2_O (90:10), at pH ~3.5. ^1^H one-dimensional data as well as ^1^H-^1^H two-dimensional TOCSY^31^ and NOESY^32^ experiments with mixing times of 80 and 150 ms were recorded at 298 K on a 700 MHz Bruker Avance III spectrometer equipped with a cryoprobe. TOCSY experiments were recorded with 8 scans for 512 increments, NOESY experiments were recorded with 32-48 scans and 512 increments depending on the signal-to-noise observed in the 1D spectra. Both TOCSY and NOESY experiments were recorded with a sweep width of 12 ppm if Trp residues were present in the peptide; otherwise a 10 ppm sweep width was used. ^1^H-^13^C as well as ^1^H-^15^N HSQC (Heteronuclear Single Quantum Coherence) spectra were also recorded at natural abundance. ^1^H-_13_C HSQC experiments were recorded with 128 scans and 256 increments with a sweep width of 10 ppm in the F2 dimension and 80 ppm in the F1 dimension. ^1^H-^15^N HSQC experiments were recorded with 256 scans and 128 increments, with a sweep width of 10 ppm in the F2 dimension and 32 ppm in the F1 dimension. For VBP-10 the 13C HSQC data were recorded in 100% D_2_O to minimize interference with the Hn-Cn resonances and the residual water signal. The data was subsequently processed using Topspin 4.0.3 (Bruker), with a solvent signal reference of 4.77 ppm at 298 K. Sequential assignment strategies^33^ were used to assign and analyze the data in the program CARA (Computer Assisted Resonance Assignment).^34^ Secondary structural features were identified by comparison of the secondary Hα shifts generated by the desired peptide to that of the equivalent values in a random coil peptide.^35^ Additional TOCSY or NOESY data at varying temperatures (288 K, 293 K, 298 K, 303 K, 308 K) were recorded to monitor temperature dependence of the amide protons.

### Structure Calculation

Inter-proton distance restraints were generated from the peak of the cross peaks present in the NOESY spectra for each peptide. TALOS-^36^ was used to predict dihedral ϕ (C^−1^-N-Cα-C) and ψ (N-Cα-C-N^+1^) backbone angles. The TALOS-N dihedral restraints and chemical shifts were also used to predict the χ^1^ (N-Cα-Cβ-Sx) and χ^2^ (Cα-Cβ-Sx-Sy) angles for the disulfide bonded Cys residues using the program Di-Sulfide and Di-Hedral prediction (DISH).^37^ Hydrogen bonds were identified via determination of backbone amide temperature coefficients. The chemical shift of the ^1^HN proton of each residue was plotted against temperature. Values > −4.6 ppb/K for the coefficient of the linear relationship were taken as indicative of a hydrogen bond being donated by the backbone amide of that particular residue.^38^ Hydrogen bond acceptors were identified through preliminary structure calculations. Initial structures (50) were calculated using the program CYANA^39^ using torsion angle simulated annealing, defining the starting coordinates and distance restraints to be used in CNS. The distance restraints generated by CYANA along with the dihedral restraints from TALOS-N and hydrogen bond restraints from temperature coefficients were used as input for the program CNS. Simulated annealing using both torsion angle and Cartesian space was conducted by CNS to generate 50 structures.^40^ These structures were then subjected to water minimization using Cartesian dynamics to generate the final structures for the peptides. Stereochemical analysis was conducted by MolProbity^41^ by comparing the generated structures to that of previously published structures. The program MOLMOL^42^ was used to display and generate images of the secondary and tertiary structure of the best 20 structures, which had good geometry, contained no violations of distances or dihedral angles above 0.2 Å or 2°, and had low energy. The structures, NMR restraints and chemical shift information have been submitted to the Protein Data Bank (PDB) and Biological Magnetic Resonance Bank (BMRB) and the accession codes are 6WQJ and 30748 for VBP-10, and 6WQL and 30749 for C2.

### Trypsin Inhibitory Assay

Trypsin inhibition was determined as previously described.^8, 43^ Synthetic peptides were dissolved and assayed for inhibitory activity in a buffer comprised of 50 mM Tris-HCl pH 7.8 and 20 mM CaCl_2_. To 20 µl of a 25 µg/mL solution of trypsin from bovine pancreas (Sigma-Aldrich), 5 µL of peptides were added to generate final concentrations ranging from 0.01 µM to 8 µM; these mixtures were pre-incubated at 37°C for 15 minutes. The reaction was initiated by the addition of 1 mM N-α-benzoyl-L-arginine-p-nitroanilide substrate (Sigma-Aldrich) and incubated for 30 min at 37°C. Soybean Bowman-Birk inhibitor (Sigma-Aldrich) was used as a positive control, and all reactions were performed in triplicate. Trypsin activity of reaction wells with no inhibitor/peptide present was designated as 100%, and the inhibitory activity of synthetic peptides was determined in relation to the no-inhibitor controls. Finally, 25 µL of 30% acetic acid were added to stop reactions, and optical absorbance was measured at 410 nm.

## ASSOCIATED CONTENT

Supporting information is available.

### AUTHOR INFORMATION

CDP peptide chemistry, NMR data analysis, structure calculations; GV and JZ assays; MF transcriptomics data analysis; RJC peptide chemistry; JSM project design and funding; KJR NMR data analysis, project design and funding. CDP wrote paper together with JSM and KJR with input from all authors.

## ACKNOWLEDGEMENTS

This work was funded by a Discovery Project from the Australian Research Council to JSM and KJR (DP190102058). CDP was supported by an University of Queensland Postgraduate Research Award.

## REFERENCES

1. Barber, C. J. S.; Pujara, P. T.; Reed, D. W.; Chiwocha, S.; Zhang, H.; Covello, P. S., The two-step biosynthesis of cyclic peptides from linear precursors in a member of the plant family Caryophyllaceae involves cyclization by a serine protease-like enzyme. J. Biol. Chem. 2013, 288 (18), 12500–12510.

2. Gillon, A. D.; Saska, I.; Jennings, C. V.; Guarino, R. F.; Craik, D. J.; Anderson, M. A., Biosynthesis of circular proteins in plants. Plant J. 2007, 53 (3), 505–515.

3. Süssmuth, R. D.; Mainz, A., Nonribosomal Peptide Synthesis—Principles and Prospects. Angew. Chem. 2017, 56 (14), 3770–3821.

4. Tobias, N. J.; Linck, A.; Bode, H. B., Natural Product Diversification Mediated by Alternative Transcriptional Starting. Angew. Chem. 2018, 57 (20), 5699–5702.

5. Mylne, J. S.; Colgrave, M. L.; Daly, N. L.; Chanson, A. H.; Elliott, A. G.; McCallum, E. J.; Jones, A.; Craik, D. J., Albumins and their processing machinery are hijacked for cyclic peptides in sunflower. Nat. Chem. Biol. 2011, 7, 257–259.

6. Elliott, A. G.; Delay, C.; Liu, H.; Phua, Z.; Rosengren, K. J.; Benfield, A. H.; Panero, J. L.; Colgrave, M. L.; Jayasena, A. S.; Dunse, K. M.; Anderson, M. A.; Schilling, E. E.; Ortiz-Barrientos, D.; Craik, D. J.; Mylne, J. S., Evolutionary origins of a bioactive peptide buried within Preproalbumin. Plant Cell 2014, 26 (3), 981–995.

7. Fisher, M. F.; Zhang, J.; Taylor, N. L.; Howard, M. J.; Berkowitz, O.; Debowski, A. W.; Behsaz, B.; Whelan, J.; Pevzner, P. A.; Mylne, J. S., A family of small, cyclic peptides buried in preproalbumin since the Eocene epoch. Plant Direct 2018, 2 (2), e00042.

8. Zhang, J.; Payne, C. D.; Pouvreau, B.; Schaefer, H.; Fisher, M. F.; Taylor, N. L.; Berkowitz, O.; Whelan, J.; Rosengren, K. J.; Mylne, J. S., An Ancient Peptide Family Buried within Vicilin Precursors. ACS Chem. Biol. 2019, 14 (5), 979–993.

9. Luckett, S.; Garcia, R. S.; Barker, J. J.; Konarev, A. V.; Shewry, P. R.; Clarke, A. R.; Brady, R. L., High-resolution structure of a potent, cyclic proteinase inhibitor from sunflower seeds J. Mol. Biol. 1999, 290 (2), 525–533.

10. Shewry, P., and Pandya, M, The 2S Albumin Storage Proteins. In Seed Proteins, Shewry, P.; Casey, R., Eds. Kluwer, Dordrecht, 1999; pp 563–586.

11. Mylne, J. S.; Hara-Nishimura, I.; Rosengren, K. J., Seed storage albumins: biosynthesis, trafficking and structures. Funct. Plant Biol. 2014, 41 (7), 671–677.

12. Jayasena, A. S.; Fisher, M. F.; Panero, J. L.; Secco, D.; Bernath-Levin, K.; Berkowitz, O.; Taylor, N. L.; Schilling, E. E.; Whelan, J.; Mylne, J. S., Stepwise Evolution of a Buried Inhibitor Peptide over 45 My. Mol. Biol. Evol. 2017, 34 (6), 1505–1516.

13. Yamada, K.; Shimada, T.; Kondo, M.; Nishimura, M.; Hara-Nishimura, I., Multiple functional proteins are produced by cleaving Asn-Gln bonds of a single precursor by vacuolar processing enzyme. J. Biol. Chem. 1999, 274 (4), 2563–2570.

14. Marcus, J. P.; Green, J. L.; Goulter, K. C.; Manners, J. M., A family of antimicrobial peptides is produced by processing of a 7S globulin protein in *Macadamia integrifolia* kernels. Plant J. 1999, 19 (6), 699–710.

15. Li, F.; Yang, X.-X.; Xia, H.-C.; Zeng, R.; Hu, W.-G.; Li, Z.; Zhang, Z.-C., Purification and characterization of Luffin P1, a ribosome-inactivating peptide from the seeds of *Luffa cylindrica*. Peptides 2003, 24 (6), 799–805.

16. Conners, R.; Konarev, A. V.; Forsyth, J.; Lovegrove, A.; Marsh, J.; Joseph-Horne, T.; Shewry, P.; Brady, R. L., Unusual Helix-Turn-Helix Protease Inhibitory Motif in a Novel Trypsin Inhibitor from Seeds of Veronica (*Veronica hederifolia L*.). J. Biol. Chem. 2007, 282 (38), 27760–27768.

17. Nolde, S. B.; Vassilevski, A. A.; Rogozhin, E. A.; Barinov, N. A.; Balashova, T. A.; Samsonova, O. V.; Baranov, Y. V.; Feofanov, A. V.; Egorov, T. A.; Arseniev, A. S.; Grishin, E. V., Disulfide-stabilized Helical Hairpin Structure and Activity of a Novel Antifungal Peptide EcAMP1 from Seeds of Barnyard Grass (*Echinochloa crus-galli*). J. Biol. Chem. 2011, 286 (28), 25145–25153.

18. Oparin, P. B.; Mineev, K. S.; Dunaevsky, Y. E.; Arseniev, A. S.; Belozersky, M. A.; Grishin, E. V.; Egorov, T. A.; Vassilevski, A. A., Buckwheat trypsin inhibitor with helical hairpin structure belongs to a new family of plant defence peptides. Biochem. J. 2012, 446 (1), 69–77.

19. Wishart, D. S.; Sykes, B. D.; Richards, F. M., Relationship between nuclear magnetic resonance chemical shift and protein secondary structure. J. Mol. Biol. 1991, 222 (2), 311–333.

20. Wishart, D. S., The chemical shift index: A fast and simple method for the assignment of protein secondary structure through NMR spectroscopy. Biochemistry 1992, 31 (6), 1647–1651.

21. Schroeder, C. I.; Rosengren, K. J., Three-Dimensional Structure Determination of Peptides Using Solution Nuclear Magnetic Resonance Spectroscopy. In Snake and Spider Toxins: Methods and Protocols, Priel, A., Ed. Springer US: New York, NY, 2020; pp 129–162.

22. Barnham, K. J.; Dyke, T. R.; Kem, W. R.; Norton, R. S., Structure of neurotoxin B-IV from the marine worm *Cerebratulus lacteus*: a helical hairpin cross-linked by disulphide bonding. J. Mol. Biol. 1997, 268 (5), 886–902.

23. Bhunia, A.; Domadia, P. N.; Torres, J.; Hallock, K. J.; Ramamoorthy, A.; Bhattacharjya, S., NMR structure of pardaxin, a pore-forming antimicrobial peptide, in lipopolysaccharide micelles: mechanism of outer membrane permeabilization. J. Biol. Chem. 2010, 285 (6), 3883–3895.

24. Möller, C.; Rahmankhah, S.; Lauer-Fields, J.; Bubis, J.; Fields, G. B.; Marí, F., A Novel Conotoxin Framework with a Helix−Loop−Helix (Cs α/α) Fold. Biochemistry 2005, 44 (49), 15986–15996.

25. Chagot, B.; Pimentel, C.; Dai, L.; Pil, J.; Tytgat, J.; Nakajima, T.; Corzo, G.; Darbon, H.; Ferrat, G., An unusual fold for potassium channel blockers: NMR structure of three toxins from the scorpion *Opisthacanthus madagascariensis*. Biochem. J. 2005, 388 (1), 263–271.

26. Srinivasan, K. N.; Sivaraja, V.; Huys, I.; Sasaki, T.; Cheng, B.; Kumar, T. K.; Sato, K.; Tytgat, J.; Yu, C.; San, B. C.; Ranganathan, S.; Bowie, H. J.; Kini, R. M.; Gopalakrishnakone, P., kappa-Hefutoxin1, a novel toxin from the scorpion *Heterometrus fulvipes* with unique structure and function. Importance of the functional diad in potassium channel selectivity. J. Biol. Chem. 2002, 277 (33), 30040–30047.

27. Shai, Y.; Fox, J.; Caratsch, C.; Shih, Y.-L.; Edwards, C.; Lazarovici, P., Sequencing and synthesis of pardaxin, a polypeptide from the Red Sea Moses sole with ionophore activity. FEBS Lett 1988, 242 (1), 161–166.

28. Tyndall, J. D. A.; Nall, T.; Fairlie, D. P., Proteases Universally Recognize Beta Strands In Their Active Sites. Chemical Reviews 2005, 105 (3), 973–1000.

29. Franke, B.; Jayasena, A. S.; Fisher, M. F.; Swedberg, J. E.; Taylor, N. L.; Mylne, J. S.; Rosengren, K. J., Diverse cyclic seed peptides in the Mexican zinnia (*Zinnia haageana*). Peptide Science 2016, 106 (6), 806–817.

30. Zhang, Y.; Lee, B.; Du, W.-X.; Lyu, S.-C.; Nadeau, K. C.; Grauke, L. J.; Zhang, Y.; Wang, S.; Fan, Y.; Yi, J.; McHugh, T. H., Identification and Characterization of a New Pecan [Carya *illinoinensis* (Wangenh.) K. Koch] Allergen, Car i 2. J. Agric. Food Chem. 2016, 64 (20), 4146–4151.

31. Braunschweiler, L.; Ernst, R. R., Coherence transfer by isotropic mixing: Application to proton correlation spectroscopy. J. Magn. Reson. 1983, 53 (3), 521–528.

32. Jeener, J.; Meier, B. H.; Bachmann, P.; Ernst, R. R., Investigation of exchange processes by two-dimensional NMR spectroscopy. J. Chem. Phys. 1979, 71 (11), 4546–4553.

33. Wüthrich, K., NMR of proteins and nucleic acids. New York: John Wiley & Sons: 1986.

34. Keller, R. L. J., The computer aided resonance assignment tutorial. 1 ed.; CANTINA Verlag: 2004.

35. Wishart, D.; Bigam, C.; Holm, A.; Hodges, R.; Sykes, B., 1 H, 13 C and 15 N random coil NMR chemical shifts of the common amino acids. I. Investigations of nearest-neighbor effects. J. Biomol. NM. 1995, 5 (1), 67–81.

36. Shen, Y.; Bax, A., Protein backbone and sidechain torsion angles predicted from NMR chemical shifts using artificial neural networks. J. Biomol. NM. 2013, 56 (3), 227–241.

37. Armstrong, D. A.; Kaas, Q.; Rosengren, K. J., Prediction of disulfide dihedral angles using chemical shifts. Chem. Sci. 2018, 9 (31), 6548–6556.

38. Cierpicki, T.; Otlewski, J., Amide proton temperature coefficients as hydrogen bond indicators in proteins. J. Biomol .NMR 2001, 21 (3), 249–261.

39. Güntert, P., Automated NMR Structure Calculation With CYANA. In Protein NMR Techniques, Downing, A. K., Ed. Humana Press: Totowa, NJ, 2004; pp 353–378.

40. Brünger, A. T.; Adams, P. D.; Clore, G. M.; Delano, W. L.; Gros, P.; Grosse-Kunstleve, R. W.; Jiang, J. S.; Kuszewski, J.; Nilges, M.; Pannu, N. S.; Read, R. J.; Rice, L. M.; Simonson, T.; Warren, G. L., Crystallography & NMR system: A new software suite for macromolecular structure determination. Acta Crystallogr., Sect. D: Biol. Crystallogr. 1998, 54, 905–921.

41. Chen, V. B.; Arendall, W. B.; Headd, J. J.; Keedy, D. A.; Immormino, R. M.; Kapral, G. J.; Murray, L. W.; Richardson, J. S.; Richardson, D. C., MolProbity: all-atom structure validation for macromolecular crystallography. Acta Crystallogr., Sect D: Biol. Crystallogr. 2010, 66, 12–21.

42. Koradi, R.; Billeter, M.; Wüthrich, K., MOLMOL: A program for display and analysis of macromolecular structures. J. Mol. Graphic. 1996, 14, 51–55.

43. James, A. M.; Jayasena, A. S.; Zhang, J.; Berkowitz, O.; Secco, D.; Knott, G. J.; Whelan, J.; Bond, C. S.; Mylne, J. S., Evidence for Ancient Origins of Bowman-Birk Inhibitors from *Selaginella moellendorffii*. Plant Cell 2017, 29 (3), 461–473.

44. Shi, T.; Li, R.; Chen, Q.; Li, Y.; Pan, F.; Chen, Q., De novo sequencing of seed transcriptome and development of genic-SSR markers in common buckwheat (*Fagopyrum esculentum*). New Strategies in Plant Improvement 2017, 37 (12), 1–15.

