## supporting information for "Defining the familial fold of the vicilin-buried peptide family"

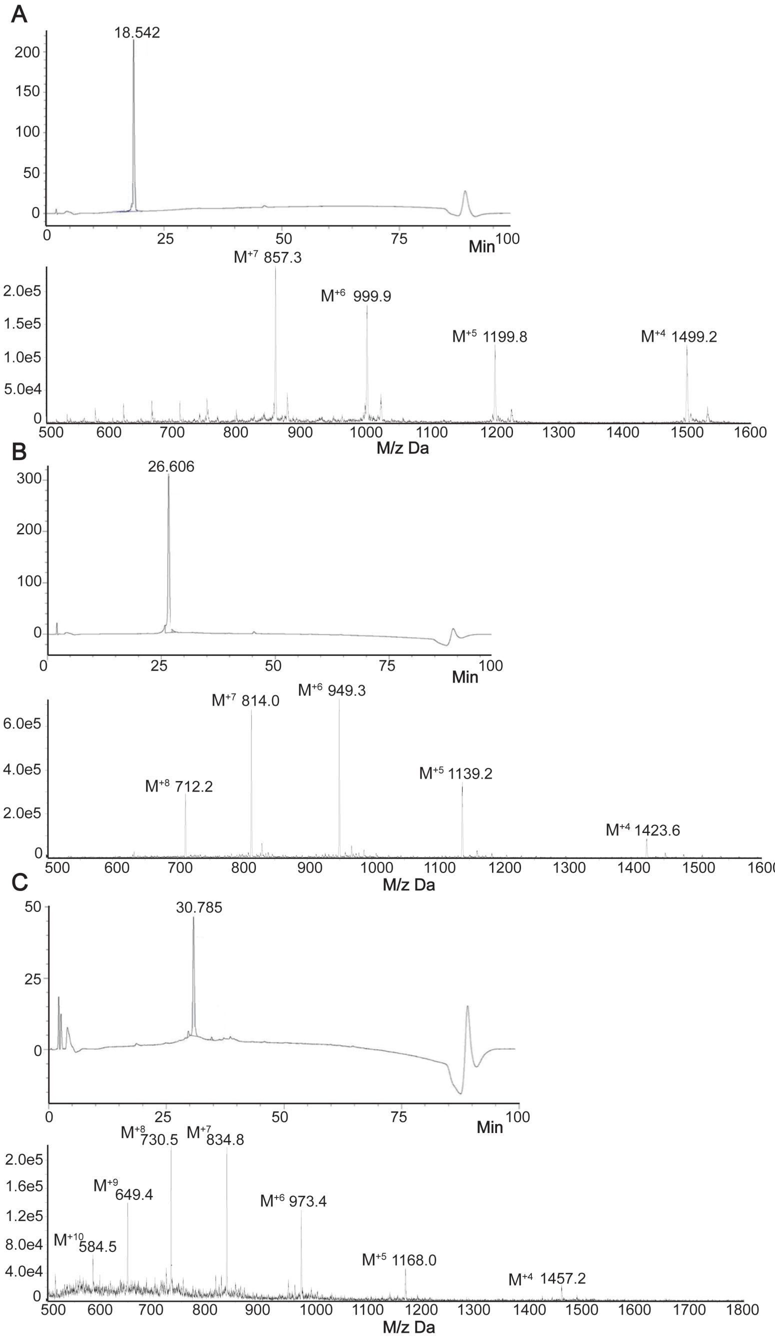


**S1 | HPLC Chromatograms showing peptide elution over time and Mass Spectra of the Synthesized VBPs.** Mass is shown in Da with M/z ratio for each peak shown. These include (A) VBP-8; (B) Luffin P1 and (C) C2.

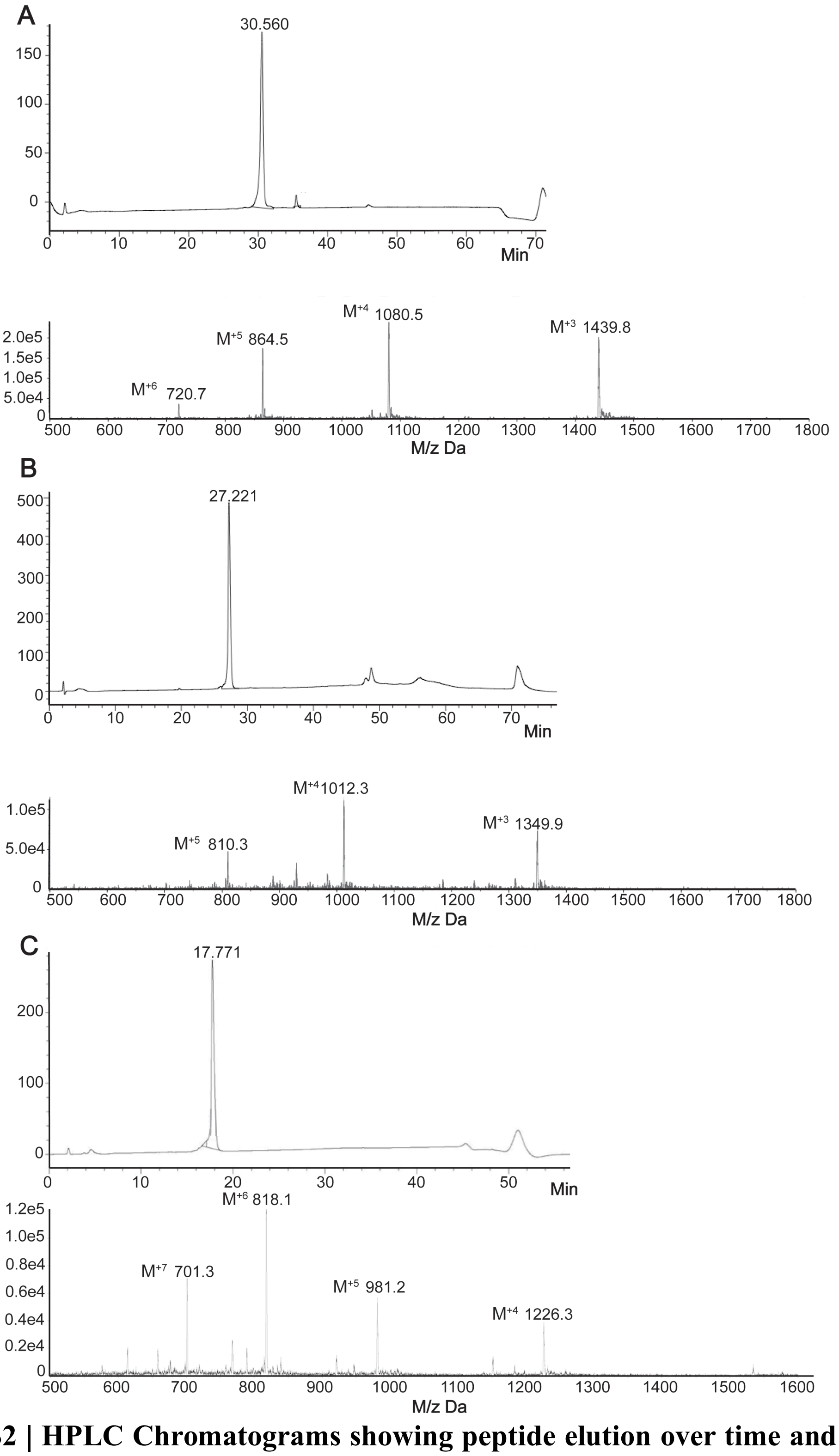


**S2 | HPLC Chromatograms showing peptide elution over time and Mass Spectra of the Synthesized VBPs.** Mass is shown in Da with *m/z* ratio for each peak shown. (A) VBP-10; (B) VhTI and (C) VBP-6.

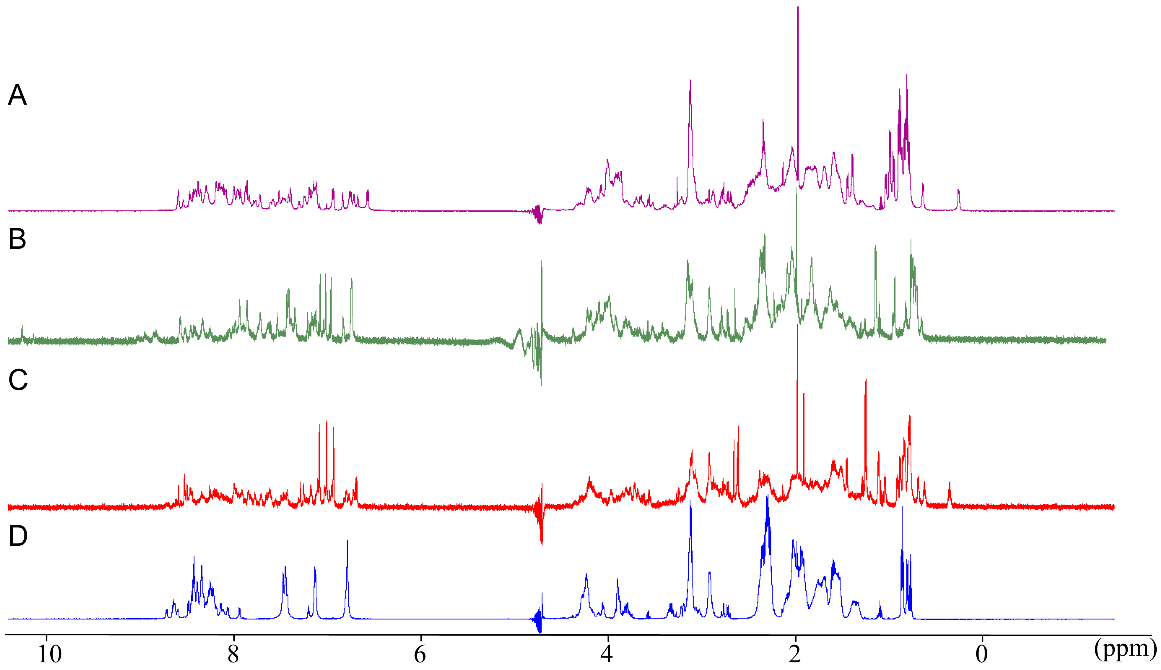


**S3 | Comparison of 1D $^1$H NMR spectra of VBPs**. (A) C2; (B) VBP-10; (C) VhTI and (D) VBP-6. The spectra show all detected $^1$H signals between 0 and 12 ppm.

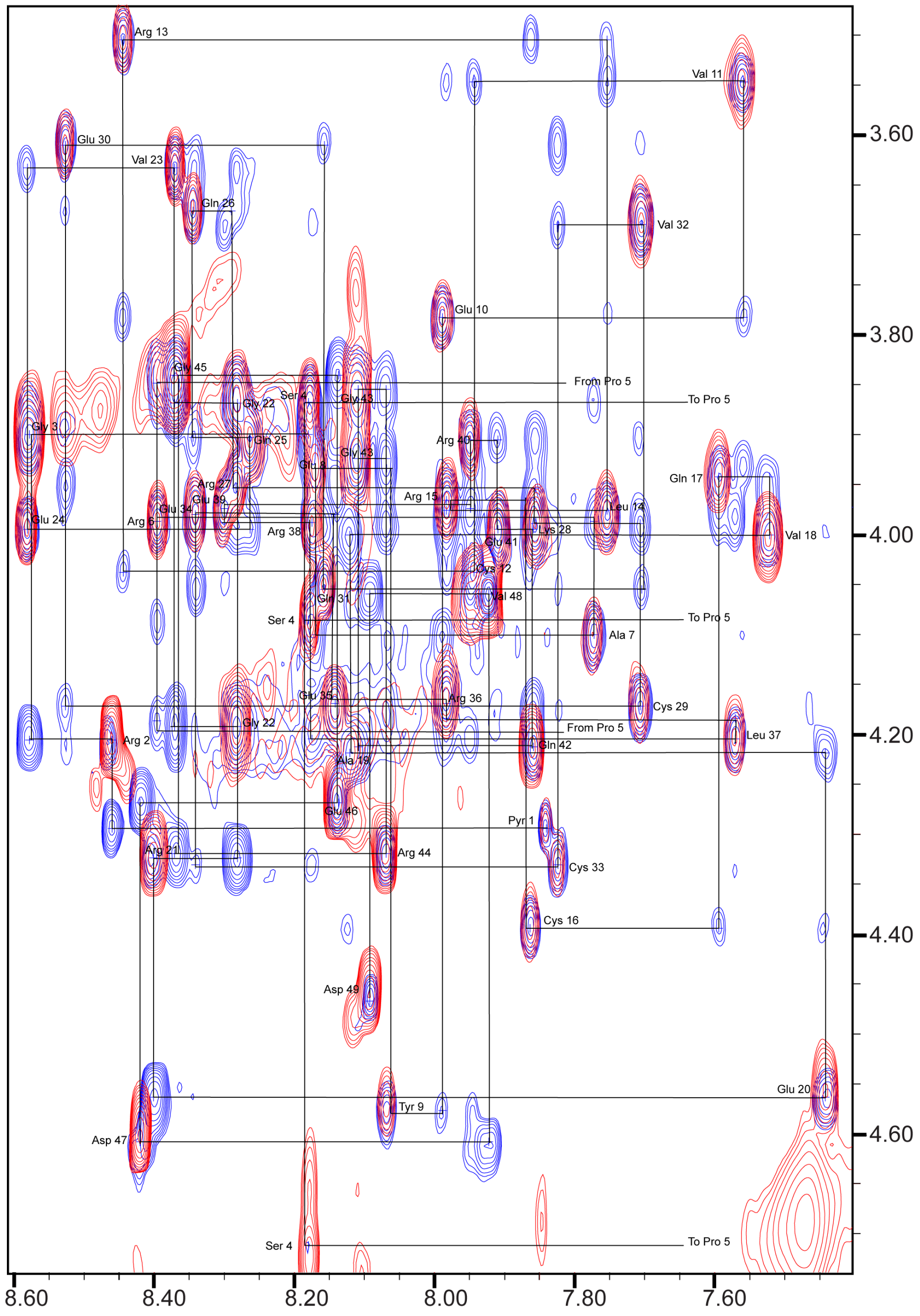


**S4 | Sequential walk assignment of the Fingerprint Region of C2**. TOCSY spectrum (80 ms mixing time) is colored red and NOESY (150 ms mixing time) spectrum colored blue. Recorded at 700 MHz and 298 K.

**a**

MVTKAKIPLFLFLSALFLALVCSSLALETEDLSNELNPHHDPESHRWEFQQCQERCQHEERGQRQAQQCQRRCEEQLRER
EREREREEIVDPREPRKQYEQCRETCEKQDPRQQPQCERRCERQFQEQQERERRERRRGRDDDDKENPRDPREQYRQCEE
HCRRQGQGQRQQQQCQSRCEERFEEEQRRQEERERRRGRDNDDEENPRDPREQYRQCQEHCRRQGQGQRQQQQCQSRCEE
RLEEEQRKQEERERRRGRDEDDQNPRDPEQRYEQCQQQCERQRRGQEQQLCRRRCEQQRQQEERERQRGRDRQDPQQQYH
RCQRRCQTQEQSPERQRQCQQRCERQYKEQQGREWGPDQASPRRESRGREEEQQRHNPYYFHSQGLRSRHESGEGEVKYL
ERFTERTELLRGIENYRVVILEANPNTFVLPYHKDAESVIVVTRGRATLTFVSQERRESFNLEYGDVIRVPAGATEYVIN
QDSNERLEMVKLLQPVNNPGQFREYYAAGAQSTESYLRVFSNDILVAALNTPRDRLERFFDQQEQREGVIIRASQEKLRA
LSQHAMSAGQRPWGRRSSGGPISLKSQRSSYSNQFGQFFEACPEEHRQLQEMDVLVNYAEIKRGAMMVPHYNSKATVVVY
VVEGTGRFEMACPHDVSSQSYEYKGRREQEEEESSTGQFQKVTARLARGDIFVIPAGHPIAITASQNENLRLVGFGINGK
NNQRNFLAGQNNIINQLEREAKELSFNMPREEIEEIFERQVESYFVPMERQSRRGQGRDHPLASILDFAGFF*

**b**

...RSTDMVHRCEEKCEDDFERQQRGGGGSSDEGNLHLELKSEKPQQELEECRNLCRSKMWSTDMVHRCENKCEEKFERQ
QRGGGSDDEERLDLELKSDEPRQELEECLDLCRTLRWSPYMVHRCEKECEDKFERQQKRGGGSDEEEDKLDPGFKSEKPQ
QELEECQNVCRMKRWSTEMVHRCEKKCEEKFERQQRGGGGSSDEGKLGLQLKSDKPQQLLEECRY...

**c**

```
                          ...RSTDMVHRCEEKCEDDFERQQRGGGGSSDEGN--------
LHLELKSEKPQQELEECRNLCRSKMWSTDMVHRCENKCEEKFERQQRGGGSDDEERLDLELKSD-
--------EPRQELEECLDLCRTLRWSPYMVHRCEKECEDKFERQQKRGGGSDEEEDKLD-----
--PGFKSEKPQQELEECQNVCRMKRWSTEMVHRCEKKCEEKFERQQRGGGGSSDEGKLGLQLKSD
--------KPQQLLEECRY...
      ** * * ****  :**    *  ******  **. *****: **.  .*
```

**S5 | Comparison of the sequences of a pecan seed storage preprovicilin and the partial sequence of the putative precursor to BWI-2c**. a) Sequence of a Class IV preprovicilin from pecan seed (*Carya illinoinensis*), UniProtKB accession B3STU4. [30] b) Translated sequence from our assembly of RNA-seq data from the NCBI Sequence Read Archive for buckwheat seed (SRA run SRR3250237)[44] representing a partial sequence of the precursor of BWI-2c. c) An alignment of repeats within the partial BWI-2c precursor.

The predicted ER targeting signal for B3STU4 is shown in pink. CXXXC motifs are shown in bold, potential AEP cleavage sites (Asp or Asn) in red and BWI-2c in blue. The brown sequence is the mature pecan vicilin Car i 2.[30] The buckwheat raw reads were assembled using CLC Genomics Workbench 20.0.3 (QIAGEN Aarhus) using default parameters, after trimming to a Q30 quality limit and a minimum read length of 50.
